# CA3 Transiently Modulates Spatial Representation in CA1

**DOI:** 10.1101/2025.01.24.634689

**Authors:** Cynthia Rais, Maxime Maheu, J. Simon Wiegert

## Abstract

Neuronal representations of the world are dynamic. A striking example is the rapid remapping of the hippocampal spatial code, which occurs even when the environment and behavior remain unchanged. CA3 input to CA1 has been shown to exert a key role in triggering synaptic plasticity in place-encoding CA1 cells — a phenomenon which could provide the cellular foundations for the remapping of the hippocampal code for space. However, how CA3 input directly contributes to place field formation and remapping of the place code in CA1 remains incompletely understood. By combining longitudinal two-photon calcium imaging of CA1 place cells with optogenetic stimulation of presynaptic CA3 neurons in mice running on a linear treadmill, we demonstrate that CA3 transiently modulates the code for allocentric space in CA1. Activation of CA3 cells both induced a small pool of new CA1 place cells and altered the pre-existing CA1 place code. The latter occurred through an unbiased shift of existing place fields specifically, and not through modifications of place field precision or a loss of place tuning in pre-existing place cells. All CA3-driven changes in the CA1 place code were transient and almost completely disappeared by the next day. Taken together, stimulation of CA3 input to CA1 cells alters CA1-encoded spatial representations, leading to a transient decrease in place field precision rather than an overrepresentation of the stimulation zone.

**HIGHLIGHTS:**

- Optogenetic CA3 stimulation with two-photon CA1 imaging in behaving mice
- Stimulation of CA3 input induces a small pool of new CA1 place cells
- Pre-existing CA1 place fields shift without loss of tuning or precision
- CA3-driven remapping is unbiased in space and largely disappears by next day

## INTRODUCTION

How abstract environmental features are represented in brain circuits remains one of the great mysteries in neuroscience. The discovery of place cells in the dorsal hippocampus approximately half a century ago^1^ provided the first answers to this question. By combining rodent models trained to navigate experimentally controlled environments with advanced recording techniques such as calcium imaging, it was later shown that individual cells in the CA1 region of the hippocampus are tuned to fixed spatial locations in the environment^1−3^. However, the factors that drive place code formation, stabilization and remapping, remain incompletely understood^4−7^.

One influential hypothesis posits that inputs from upstream regions play a central role in the regulation of the spatial code in CA1^8,9^. The vast majority (∼84%^10^) of these inputs are mono-synaptic and originate from CA3^8−10^ with approximately half of these inputs arising from the contralateral CA3 through so-called Schaffer commissurals in rodents. Recent work has emphasized behavioral timescale plasticity (BTSP) as a key synaptic mechanism through which inputs to CA1 shape the hippocampal representation of space^14–16^. Under this framework, synaptic inputs from CA3 are strengthened when they arrive in close temporal proximity to plateau depolarization in CA1 pyramidal cells^15,17–19^.

However, these studies relied on direct postsynaptic depolarization of CA1 neurons, either through current injection or optogenetic stimulation, thereby leaving the contribution of synaptic inputs to place field formation and place code regulation unresolved. In addition, it was shown that optogenetic silencing of CA3 inputs to CA1 impairs place field precision^20^, highlighting the role of CA3 synaptic inputs in shaping the CA1 place code. Finally, several previous studies have reported only limited place cell induction upon direct CA1 activation^21–23^. Taken together, these findings support the view that direct synaptic inputs from CA3 constitute a decisive factor in the formation and refinement of spatial representation in CA1.

While previous studies have established a role for CA3 in regulating CA1 place field firing^14,18,20^, likely via synaptic plasticity mechanisms in CA1 (e.g., BTSP)^15,18^, it still remains incompletely understood to what extent and *how* CA3 inputs modify spatial information encoded in CA1. Here, we sought to determine the extent to which activation of CA3 cells alters the spatial code in CA1. Specifically, our objectives were (1) to discriminate between several potential modulation scenarios, including the de-novo emergence of place fields and the remapping of pre-existing ones, and (2) to characterize the temporal dynamics of these changes by comparing them to the spontaneous remapping naturally occurring in the hippocampus.

We combined longitudinal two-photon calcium imaging of CA1 place cells with optogenetic stimulation of CA3 neurons in awake mice, navigating on a linear track. Given that half of CA3 inputs to CA1 originate from the contralateral hippocampus^24–27^, we could conveniently manipulate CA3 activity in one hemisphere (left) while monitoring changes in CA1 spatial representation in the other hemisphere (right)^16^.

Consistent with the role of CA3 inputs in CA1 place code regulation, we observed that activation of CA3 cells modified CA1 place representation both by creating a small pool of new, distributed place cells and by shifting pre-existing place fields to new, unbiased locations; both effects also existed, albeit at reduced rate, under sham optogenetic stimulation. All induced changes of the CA1 place code were transient and did not persist until the next day, highlighting the dynamic nature of hippocampal circuits, whereby transient modifications can occur without compromising the integrity of long-term spatial coding.

## RESULTS

### Spatial navigation task

Head-fixed, water-scheduled adult mice were trained to collect a water drop by licking at a fixed location along a 2-meter-long linear treadmill divided into five distinct tactile cue sections (**Figure 1**a). Mice learned to orient in this one-dimensional space and to locate the water reward, as indicated by two behavioral features: mice slowed down specifically upon approaching the reward zone and exhibited anticipatory and zone-specific licking (**Figure 1**b).

**Figure 1.**
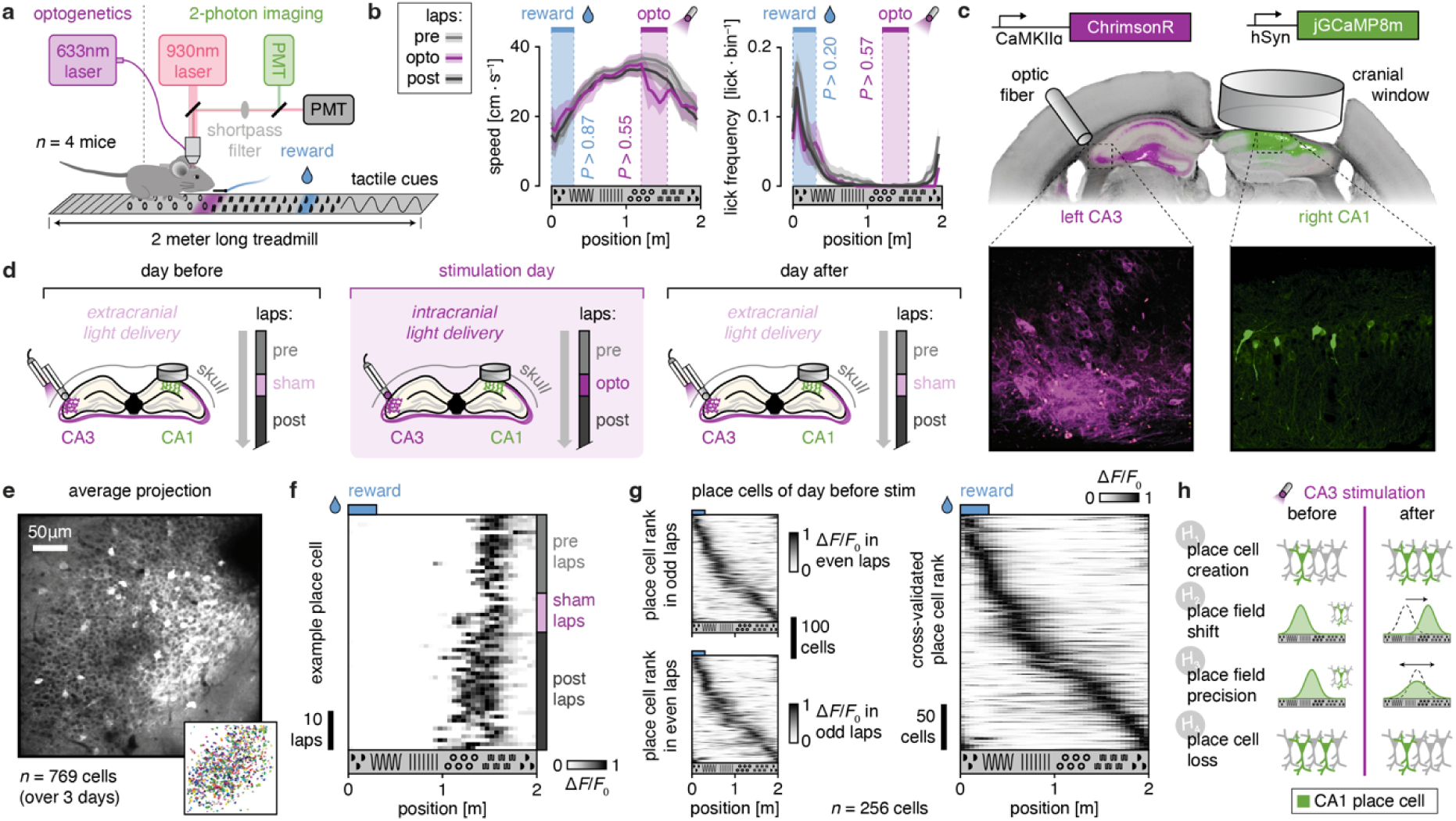
Monitoring of CA1 place cells in mice engaged in spatial navigation. **a**, Schematic depicting a head-fixed, water-scheduled mouse performing the spatial navigation task under the two-photon microscope. Both reward and optogenetic stimulation light are delivered at a fixed location on the treadmill. **b**, Mean ± s.e.m. (*n* = 4 mice) speed (left panel) and lick frequency (right panel) as a function of position on the treadmill for laps before (pre), during (opto) and after (post) optogenetic stimulation of CA3 (see panel c). Blue and magenta shaded areas denote reward delivery and optogenetic stimulation (opto) zones, respectively. No significant differences in speed and lick frequency across lap categories (pre/opto/post) for reward or opto zones. *P*-values are obtained from a Kruskal-Wallis test. **c**, Histology showing the expression of ChrimsonR (magenta; left) in left CA3 and jGCaMP8m in right CA1 (green; right). **d**, Experimental timeline. On the stimulation day (middle), optogenetic stimulation light was delivered in laps #21 to #30. On the days before and after the stimulation day, light was delivered extracranially (referred hereafter to as sham stimulation) as a control. **e**, Average projection of 2-photon time series of jGCaMP8m fluorescence in CA1 from an example mouse and corresponding cell masks identified by *Suite2p* across all 3 days (bottom right). **f**, Activity (normalized Δ*F*/*F*_0_) of an example place cell across treadmill positions (40 bins) for successive laps (from top to bottom) on the day before optogenetic manipulations. The place field of this cell was centered around ∼1.5m. **g**, Average place tuning curves across all laps (right) for all place cells identified on the day before optogenetic manipulations. Place cell rank is cross validated by analyzing separately odd and even laps (left). **h**, Hypothetical scenarios through which CA3 optogenetic stimulation could alter the pre-existing CA1 place code.

### Combined two-photon imaging and optogenetic stimulation

In mice, commissural fibers originating from CA3 make up nearly half of synaptic inputs to CA1^10,13^. Thus, we could combine simultaneous imaging of CA1 neuronal activity in one hemisphere (right CA1) with optogenetic stimulation of presynaptic CA3 neurons from the other hemisphere (left CA3) in the same animals. To monitor CA1 place cell activity, we densely labeled CA1 pyramidal neurons with a non-conditional calcium indicator (jGCaMP8m) and implanted a chronic hippocampal imaging window. To activate inputs to CA1 cells, we expressed a red light-activated soma-targeted excitatory opsin (st-ChrimsonR) in the contralateral CA3 and triggered action potentials with 633nm light delivered through an optic fiber (**Figure 1**c). With the goal of comparing the effects of CA3 optogenetic stimulation to baseline CA1 activity and to evaluate the stability of modifications induced by CA3 stimulation, we recorded CA1 activity on three consecutive days: one day before, during, and one day after CA3 stimulation (**Figure 1**d). Each recording session was divided into three consecutive blocks: 20 baseline laps without light delivery (pre laps), 10 stimulation laps, followed by a minimum of 20 laps without light delivery (post laps). We habituated mice to light stimulation patterns by delivering extracranial laser light (hereafter referred to as sham stimulation) during training sessions as well as in days before and after optogenetic stimulation (**Figure 1**b). This had the dual benefit of (1) reducing off-target effects of optogenetic light delivery on behavior and place representation, and (2) revealing the specific contribution of CA3 optogenetic activation beyond the added sensory landmark associated with optogenetic light delivery.

### Effect of optogenetic manipulations on behavior

We confirmed the absence of non-specific effects of optogenetic stimulation on behavior by comparing locomotion speed and lick rates in laps with vs. without optogenetic stimulation on the stimulation day. A small, but non-significant, deceleration was found during optogenetic stimulation (‘opto’; **Figure 1**b). Importantly, this small optogenetically induced change in running behavior remained confined to laps with optogenetic stimulation and vanished entirely in post stimulation laps. Similarly, licking behavior remained unaffected by optogenetic stimulation of CA3. Thus, optogenetic stimulation of CA3 did not significantly alter task-related behavior, allowing a direct comparison between post and pre laps.

### Identification of place cells from CA1 activity

We recorded cellular CA1 calcium activity (**Figure 1**e, **Supplementary Figure 1**) and identified place cells^1−3^: i.e., cells encoding the same location on the belt across laps (**Figure 1**f). In the day before optogenetic stimulation, we could detect 256 place cells (equivalent to 8.8 ± 2.4% of all recorded cells for each mouse; see **Methods** for place cell identification). As expected, place cells covered the entire one-dimensional space of the linear treadmill. For visualization, place cells were ranked by the center of their place field (i.e., position with the strongest associated Δ*F*/*F*_0_). More specifically, place tuning curves from odd laps were ranked based on the place field center identified from even laps (and conversely on odd laps; **Figure 1**g, left panel). Thus, averaging cell ranks obtained separately on odd and even laps gives rise to cross validated cell × position matrices (**Figure 1**g, right panel) in which the diagonal pattern is not a mere consequence of peak alignment (see **Methods** for 2-fold, odd/even lap-based cross validation). Thus, our experimental approach can detect place cells which reliably encode spatial locations across laps.

### Hypothetical modifications of CA1 place code by CA3 optogenetic activation

We reasoned that driving synaptic input to CA1 cells via CA3 optogenetic stimulation could influence place representation through 4 different, possibly co-existing scenarios (**Figure 1**h). A simple possibility is that CA3 triggers the creation of new place cells by increasing excitatory synaptic input to CA1 cells and possibly triggering plasticity mechanisms (*H*_1_; either through de-novo place cell creation or place cell unmasking^12−15^). Alternatively, the additional synaptic input created by CA3 optogenetic stimulation could alter the place field of pre-existing place cells: the place field might shift to a new location (*H*_2_; for instance towards the optogenetically stimulated zone^29^), or the precision of place fields could be altered (*H*_3_; either becoming more focused or less precise). In an extreme scenario, CA3 stimulation could even lead to a complete loss of place tuning in pre-existing place cells (*H*_4_). All these possibilities (*H*_1-4_) could occur through synaptic plasticity mechanisms acting directly at CA1 pyramidal neurons (e.g., BTSP), indirectly by recruiting local CA1 interneurons^21,30^, or via other cellular mechanisms.

### Remapping of CA1 place code

Before investigating the four hypothetical scenarios outlined above, we first assessed natural dynamics of the CA1 spatial code over time. Previous studies showed that remapping occurred naturally over time ^4−7^, which involves synaptic plasticity of CA3-inputs to CA1 cells^31^. Our chronic imaging approach enabled us to identify the same CA1 cells across days and to track their spatial tuning (**Figure 2**a,b). We focused on single-mode place fields, as this corresponded to the vast majority of place cells we could identify (**Figure 1)**, in line with previous reports^29,32^. To visualize the conservation vs. modification of place tuning across days, we identified place cells separately on three days (with cross validated ranks, as in **Figure 1**) and tracked their place tuning curves on the other two days (**Figure 2**b). This analysis revealed instability in place representation across days, with place cells rarely preserving their place tuning across days. As a control, place tuning was plotted on the same day from which ranks were obtained (using cross validation; see **Supplementary Figure 2** for cross-validated maps before averaging across odd and even laps), which showed that place representation was stable within each single day. Thus, modification of the place code across days was not a mere consequence of the way analysis was conducted and reflected a genuine instability in spatial representation across days, confirming previous findings in CA1 at this temporal scale^5^.

**Figure 2.**
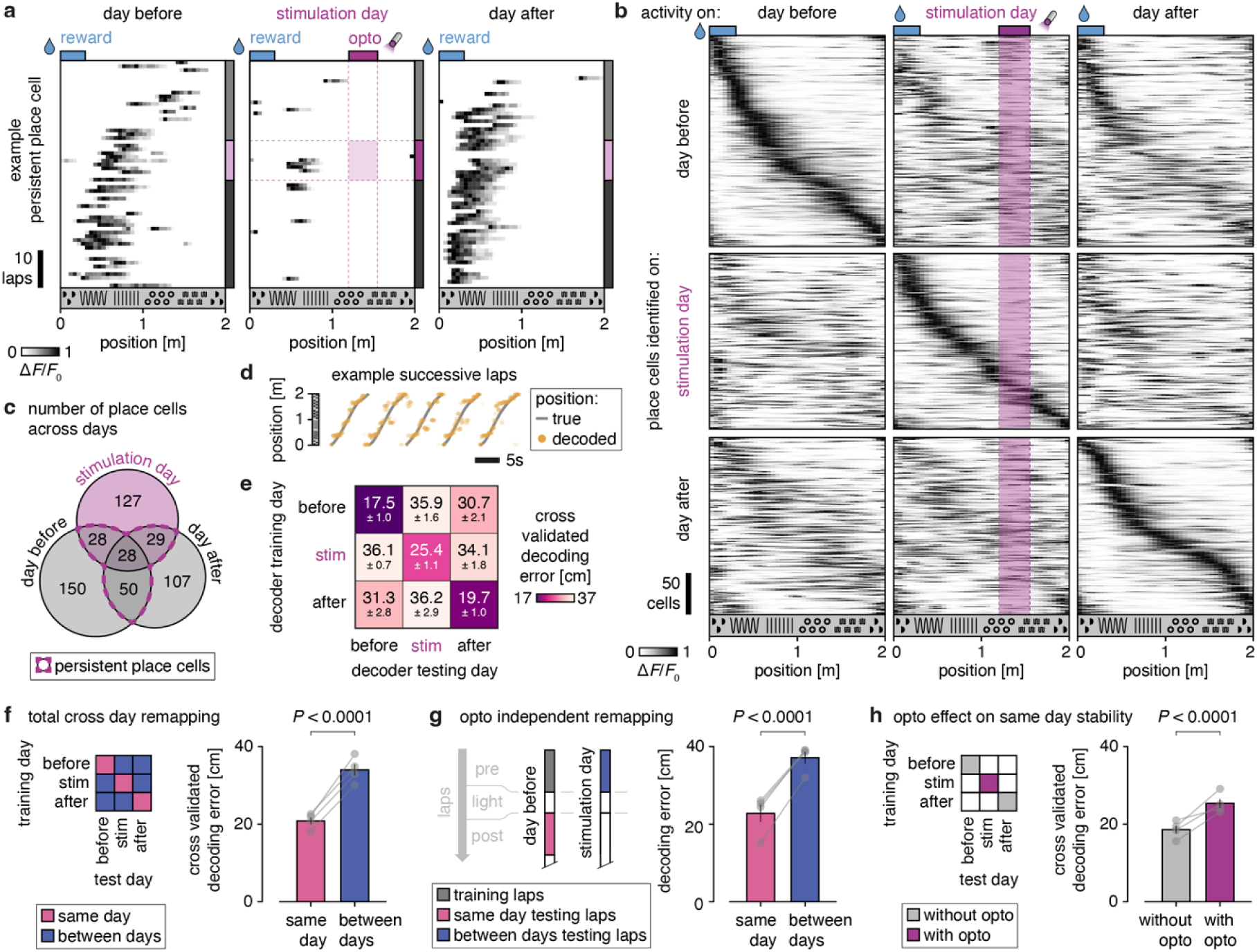
Remapping of the place code in CA1. **a**, Example of a place cell remapping across days. **b**, Place tuning curves computed separately on each day (columns) for place cells identified on a given day (rows). **c**, Number and identity of place cells for each of the 3 recording days. Persistent place cells are cells encoding space at least in two consecutive days. **d**, True animal position and position decoded from the place cell activity in an example snapshot of data (5 consecutive laps). **e**, Mean ± s.e.m. error in position decoding on all three recording days (columns) using a decoder trained on place cell activity from the same (diagonal) or different (off diagonal) day. **f**, Mean ± s.e.m. same-day vs. between-days decoding error, corresponding respectively to diagonal vs. off-diagonal cells in the matrix of panel e. **g**, Mean ± s.e.m. same-day vs. between-days decoding error before optogenetic manipulations occurred. In both cases, pre laps from the day before stimulation were used to train the decoder. Same-day decoding error was estimated by using post laps from the day before stimulation, while between-days decoding error was estimated using pre laps from the stimulation day. **h**, Mean ± s.e.m. same-day decoding error separately for days without (day before and after) vs. with optogenetic stimulation. In all panels, s.e.m. was computed across mice (*n* = 4) and *P*-values are obtained from permutation tests.

Next, we quantified the extent of remapping more directly by training a decoder^33^ to use place cell activity from one of the three recording days to decode animal position on the other days (**Figure 2**d-e; see **Methods**). As control, we trained and evaluated decoder performance using data from the same day. To ensure a fair comparison, we used a two-fold cross validation scheme where odd and even laps of the same day served as distinct and counter-balanced training and testing sets (see **Methods**). In this analysis, remapping is directly reflected by the absolute distance between the true position and the most likely position as estimated by the decoder (hereafter referred to as decoding error). Using the decoder, we confirmed remapping across days, which was indicated by a larger decoding error when the decoder was trained and tested on different days vs. on the same day (**Figure 2**f). Notably, remapping also occurred independently of optogenetic manipulations, as a decoder trained on pre laps of the day before stimulation generalized better to post laps of the same day than to pre laps of the stimulation day (i.e., all laps occurring before optogenetic stimulation; **Figure 2**g).

### Optogenetic stimulation transiently reduced the stability of the CA1 place code

Next, we asked whether optogenetic stimulation of CA3 would modulate spatial tuning of CA1 place cells. Despite spontaneous remapping across days, we observed that optogenetic stimulation of CA3 increased the decoding error on the same day (stimulation day) compared to the other days (**Figure 2**h). Thus, similar to acute optogenetic silencing^20^, stimulation of CA3 cells also acutely reduced the stability of the place code in CA1. We aimed to identify the cause of this optogenetically-induced reduction of stability, addressing each of the four hypotheses presented above (**Figure 1**), starting with the optogenetically-induced formation of novel place cells (*H*_1_).

### Induction of novel CA1 place cells by CA3 optogenetic stimulation

To identify CA1 place cells induced by CA3 optogenetic stimulation (*H*_1_), we sorted CA1 cells on the stimulation day using a two-factor scheme: On the one hand, cells were classified according to their response to the optogenetic stimulation as non-/responding, on the other hand, they were categorized based on their spatial tuning distinguishing neurons with a pre-existing place field, an induced place field, or no place field (**Figure 3**a, b: see **Methods**). Responding CA1 cells represented 18% (698 / 3784) of the recorded cells on the stimulation day (**Figure 3**c). Induced place cells were defined as cells encoding position specifically in post- but not in pre-stimulation laps. On the day with optogenetic stimulation, 44% (94 / 212) of all place cells were induced (**Figure 3**d). Having incorporated sham light stimulation in the design, we could control for place cells appearing spontaneously or induced by perception of the laser light, which in itself provides an additional visual input to the animal^34^. Indeed, also on the day before optogenetic stimulation place cells were induced by the laser light alone, albeit at a lower rate, resulting in 30% (77 / 256) induced place cells. Importantly, the higher induction rate observed during CA3 optogenetic stimulation indicates that activation of CA3 inputs specifically enhances the formation of novel place cells beyond the effects of light exposure alone. Interestingly, only 13% (12 / 94) of the optogenetically induced place cells were directly responding to CA3 optogenetic stimulation (dashed circle in **Figure 3**d). Rather, the optogenetically induced place cells tended to appear during laps following optogenetic stimulation and not during the optogenetic stimulation laps where the rate of new place cell formation was similar on all 3 days (**Figure 3**e).

**Figure 3.**
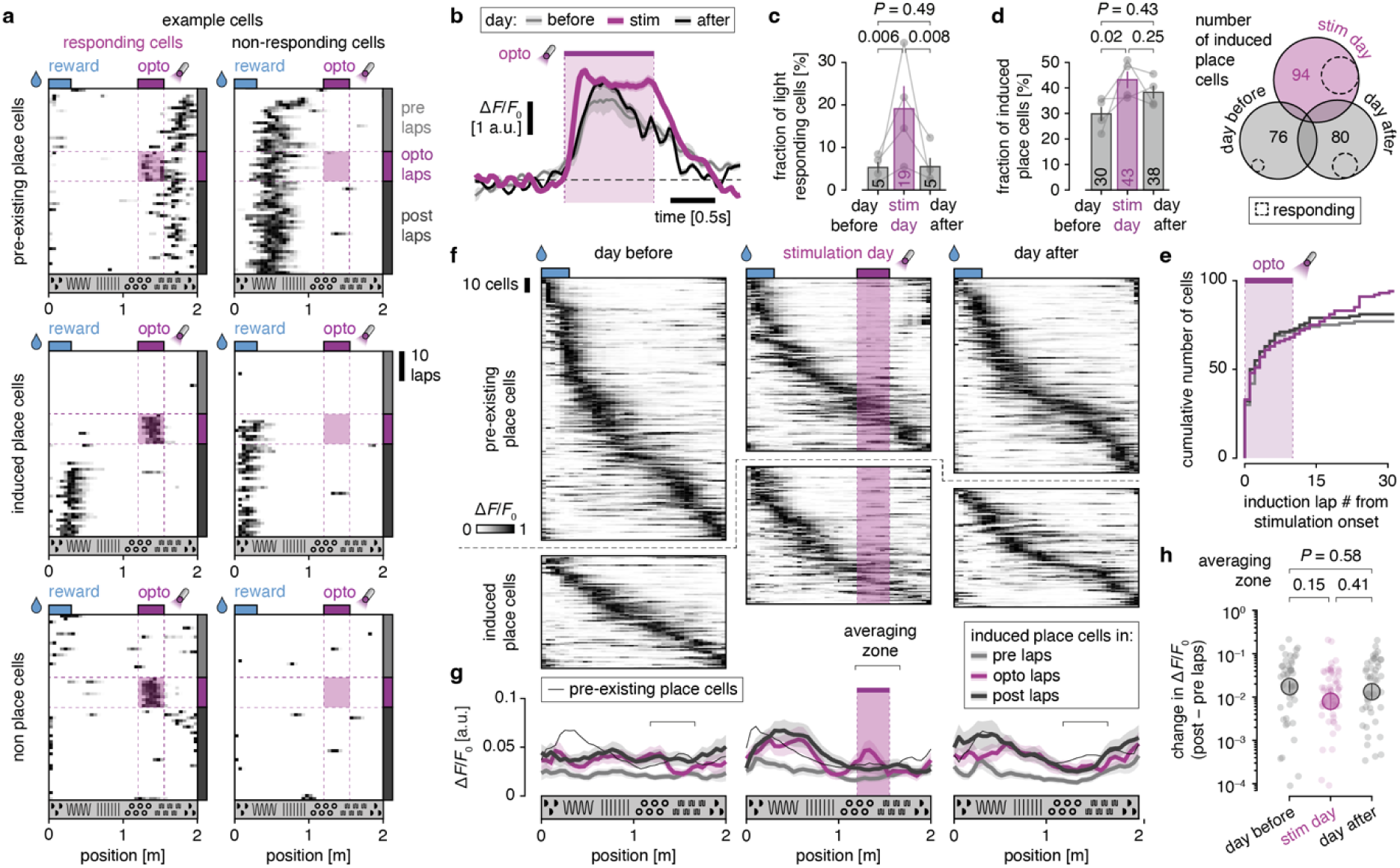
Induction of place cells in CA1 by CA3 optogenetic stimulation. **a**, Matrix of representative examples of CA1 cells encoding position (rows; pre-existing/induced/non-place cells) and/or responding to CA3 optogenetic stimulation (columns; responding/non-responding cells). **b**, Mean ± s.e.m. activity of CA1 responding cells evoked by CA3 optogenetic stimulation (purple; *n* = 698 cells) and by sham stimulation-stimulation on the day before (grey, *n* = 236) and the day after (black, *n* = 282). **c**, Mean ± s.e.m. (*n* = 4 mice) fraction of CA1 cells characterized as responding to sham (days before and after) and optogenetic stimulation. P-values are obtained from a permutation test. **d**, Mean ± s.e.m. fraction of induced place cells (*n* = 4 mice) and corresponding Venn diagram representing the number of place cells induced by sham and/or optogenetic stimulation. Induced place cells are cells encoding position in post but not in pre laps (Fisher exact test on the proportion of induced place cells among all recorded place cells). **e**, Cumulative count of induced place cells as a function of the lap number on which they appear (relative to the first lap with optogenetic stimulation; shaded area indicates the 10 laps with optogenetic stimulation). **f**, Separate place tuning curves for pre-existing (top) and induced (bottom) place cells on three days (columns), with cross validated cell ranks. Pre-existing place cells are cells encoding position already in pre-stimulation laps. **g**, Average activity of pre-existing and induced place cells (separately for pre/opto/laps) along treadmill position. **h**, Mean difference in activity ± s.e.m. (across cells) between post and pre laps for induced place cells in the zone. Shaded dots are individual cells. *P*-values are obtained from an independent Student *t*-test.

We then sought to determine the place preference of these induced place cells, considering the possibility that induced place cells could exhibit place fields centered on locations other than the fixed location of the optogenetic stimulation^22−24^, in line with BTSP where synaptic inputs active before or after an instructive signal may get strengthened. Compared to pre-existing place cells, induced place cells were not preferentially located on any specific treadmill location (**Figure 3**f). To quantify the change in position-resolved place cell activity between post and pre laps, we defined a spatial zone encompassing the part of the belt which included the optogenetic stimulation zone (**Figure 3**g). Place cell activity was larger in post than pre laps, a simple consequence of the way induced place cells are defined (i.e., cells with place tuning in post but not in pre laps). However, this increase in post lap activity could also be observed on the day before (**Figure 3**h), precluding the possibility of a CA3-optogenetically induced bias in place tuning. Thus, consistent with *H*_1_, optogenetic activation of CA3 triggered the formation of a small pool of novel place cells, which were distributed along the linear track without any apparent bias. However, the stimulation of place cells was not directly associated with responses during the optogenetic stimulation or the location of the stimulation. Thus, CA3 stimulation alone is not sufficient to reliably bias the formation of novel place fields.

### Perturbation of pre-existing CA1 population place code by CA3 optogenetic stimulation

Given the relatively small amount of place cells induced via CA3 stimulation (especially relative to sham stimulation), we next tested the possibility that CA3 inputs to CA1 could alter the pre-existing place code more generally (*H*_2-4_). We assessed whether the spatial information carried at the population level in pre stimulation laps was preserved in post stimulation laps. We reasoned that if this was the case, a decoder trained on the pre stimulation lap population code should be able to decode the animal position in post stimulation laps (see **Methods**, **Figure 4**a). To ensure that our conclusions only reflect modifications of the pre-existing place code, we focused exclusively on pre-existing place cells (i.e., excluding induced place cells). We first created decoding matrices depicting probability distributions over positions (as estimated by the decoder) conditionally on the true position of the animal (**Figure 4**b). A clear diagonal pattern was observed on all three days demonstrating that at least part of the pre-existing place code in pre laps was conserved in post laps. We then compared the conservation of the place code between pre and post laps across days and observed a larger decoding error (see **Methods**) on the day of optogenetic stimulation compared to days without simulation. This effect was present at a large fraction of the track (**Figure 4**c), thus resulting in a larger overall decoding error on the stimulation day (**Figure 4**d). Therefore, CA3 optogenetic activation alters the pre-existing population place code in CA1.

**Figure 4.**
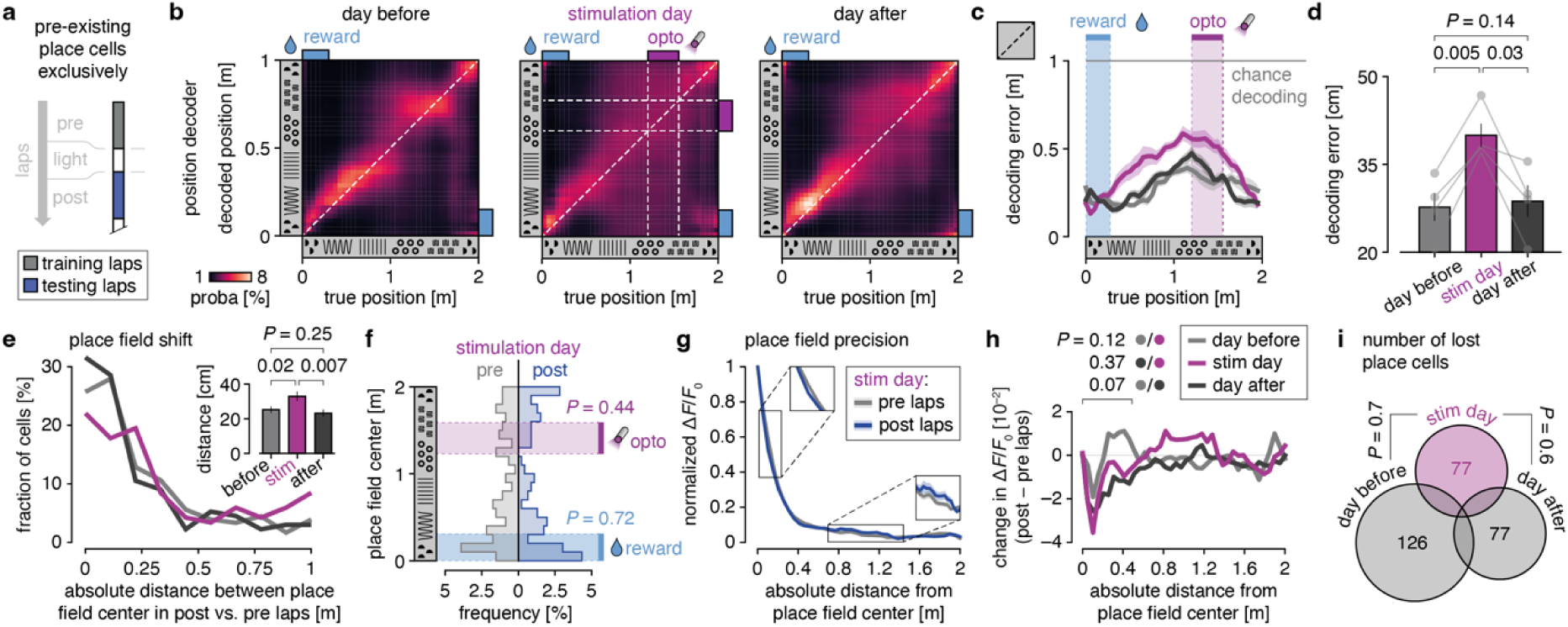
CA3-driven alteration of pre-existing CA1 place code through place field shifting. **a**, Perturbation of pre-existing population place code is assessed by comparing the activity of pre-existing place cells exclusively (as opposed to induced place cells) in post vs. pre laps. **b**, Probability of animal position detected by a decoder trained on pre laps vs. true position in post laps on the 3 consecutive experiment days. **c**, Mean ± s.e.m. (*n* = 4 mice) decoding error of the true position of the animal (corresponding to diagonals in panel b) on the 3 days. **d**, Mean ± s.e.m. (*n* = 4 mice) decoding error averaged across the entire treadmill length (in panel c). *P*-values are obtained from a permutation test. **e**, Inverse cumulative distribution of the shift in the center of the place field from pre to post laps on all 3 days. Inset depicts the average place field shift on each day. *P*-values are obtained from a permutation test. **f**, Histogram of place field center separately on pre and post laps of the stimulation day. *P*-values are obtained from a Fisher exact test. **g**, Mean ± s.e.m. (*n* = 118 cells) normalized activity aligned to the center of the place field separately in pre and post laps of the stimulation day. **h**, Change in activity between post and pre laps (difference between curves in panel g) on all 3 days as a function of the absolute distance to the center of the place field. **i**, Venn diagram representing the number of lost place cells on all 3 days. Lost place cells are cells encoding position in pre but not in post laps. *P*-values are obtained from a Fisher exact test.

### CA3-driven perturbation of pre-existing place code is caused specifically by a shift of place fields

We next aimed to identify the precise modifications of the spatial code that were induced by optogenetic CA3 stimulation. As mentioned above, CA3 inputs could shift the place field of pre-existing place cells to new locations (*H*_2_) or modulate their precision (*H*_3_) potentially leading to a complete loss of place tuning (*H*_4_).

To measure the extent of place field shift (*H*_2_), we computed the absolute distance between the peak position of place fields in post vs. pre laps (**Figure 4**e). We found a ∼9 cm larger (equivalent to 23% of a tactile cue section, or 5% of the entire linear track) place field shift between post and pre laps specifically on the stimulation day compared to the other two days without optogenetic manipulations. Interestingly, place fields shifted along the linear track with no clear preference (compared to pre laps; **Figure 4**f) akin to the unbiased distribution of induced place cells (**Figure 3**). In particular, the center of place fields shifted in post laps did not cluster near the optogenetic stimulation zone.

One alternative explanation is that place fields simply got less precise and noisier in post laps (*H*_3_), thus explaining why the center of the place field is more subject to undirected shift. To address this possibility, we expressed place cells’ normalized activity as a function of the absolute distance to the center of the place field separately in pre and post laps. In this analysis, a sharp decay in normalized activity indicates a precise place field. On the stimulation day, place field precision was strikingly similar in pre and post laps (**Figure 4**g). The only change we could detect is a slight increase in precision in post laps, which was visible on all 3 days (see negative peak at ∼10cm in **Figure 4**h), and likely reflected a progressive refinement of the neural code throughout the session. Critically, no pre-to-post modifications in place field precision specifically occurred on the stimulation day, precluding *H*_3_ and supporting the idea that place fields are shifted (*H*_2_), rather than broadened upon CA3 stimulation.

Finally, we considered the possibility that the optogenetic stimulation of CA3 could have led to an increased loss of place tuning in some of the pre-existing place cells (*H*_4_). Contrary to this hypothesis, the number of lost place cells between pre and post stimulation laps was comparable on all 3 days and did not specifically increase on the stimulation day (**Figure 4**i). Thus, the alteration of the pre-existing CA1 place code by CA3 optogenetic stimulation was caused specifically by a shift of place fields and not by a change of place field precision or a complete loss of place tuning.

## DISCUSSION

While it is now widely acknowledged that individual cells in the hippocampus encode specific environmental features such as fixed spatial locations, it remains incompletely understood how these representations form and evolve over time. In particular, the synaptic mechanisms driving place cell formation and place field shifts still remain elusive. While BTSP has emerged as a mechanism that can explain the temporal dynamics of place field formation, the precise role of CA3 synaptic inputs in forming and regulating place fields is still a matter of investigation. How does synaptic input from CA3 itself contribute to place field formation and precision? Is CA3-mediated input sufficient to drive postsynaptic depolarization and subsequent synaptic potentiation as suggested previously^18^? Here, we tested the extent to which CA3 inputs to CA1 play a role in shaping CA1 place representation. We show that optogenetic stimulation of CA3 neurons in the left hemisphere increased the propensity of *de novo* place cell formation in CA1 in the right hemisphere with unbiased place field locations. In addition, we show that this manipulation leads to a transient perturbance of the pre-existing population place code. This perturbation was underpinned specifically by a shift of existing place fields to new, unbiased locations and not by a reduction of place field precision or the complete loss of pre-existing place cells.

Although CA3 optogenetic stimulation favored the creation of novel place cells, these place cells did often not map to the location of stimulation. In addition, many cells responding to the optogenetic stimulation did not become place cells. This is in line with previous studies reporting variable success rates in the number, specificity and stability of induced place cells including following direct CA1 stimulation^14,15,18,21–23,36^. One possibility is that the CA3-driven synaptic input to CA1 evoked in our experiments was insufficient to induce plateau potentials, which are considered to be essential for robust place cell formation^14–16,36,37^. Rather, successful induction of place fields may require additional conditions, such as temporally coordinated synaptic input to distal and proximal CA1 dendrites from layer 3 neurons of entorhinal cortex (EC3) and CA3 neurons, respectively. This coordinated synaptic drive has been shown to be critical for triggering dendritic plateau potentials that underlie place cell emergence^14^. In addition, the strong local inhibitory activity that characterizes hippocampal network activity^17,18,32−35^ provides only a limited window for place field induction^23,40,42^. The location of newly formed place cells and the remapping of existing place cells after optogenetic CA3 stimulation may therefore be determined in large part by these additional factors.

Aside from de-novo place cell induction, we observed an acute shift of the place code in CA1 after optogenetic stimulation of CA3, which was restricted to post-stimulation laps of the stimulation day and disappeared on the next day. Similar to place cell induction, this suggests that a phasic change in CA3 inputs to CA1 alone is not sufficient to profoundly alter the existing neural code on a long-time scale. Thus, the long-term modification of hippocampal spatial representation likely relies on a joint combination of synaptic inputs along with other cellular mechanisms such as the ones described above for place cell induction. Phase-locking of CA3 activity to hippocampal oscillations, especially in the theta frequency range constitutes another important factor for place cell modulation^14,43^. Additionally, long-range afferents, including noradrenergic input from the locus coeruleus^44,45^ or glutamatergic input from the medial septum^46^ could also possibly play a key role in triggering long-term modifications of CA1 place representation. Notably, BTSP may still underlie the shift of place cells we observed: considering the protracted window that is permissive for the integration of synaptic inputs into place fields, additional, strong synaptic input provided by our optogenetic stimulation may have opened a plasticity window enabling the shift of a place field to a position where natural, spatially tuned synaptic inputs were potentiated.

Another reason for our limited ability to reliably drive strong responses in CA3 cells may reside in our experimental design: Although CA3 provides the vast majority of CA1 inputs^9,10,25−27^, our optogenetic stimulation was limited to the somata of contralateral CA3 cells, thus, triggering at most 50% of all available CA3 inputs to CA1 cells via direct, mono-synaptic connections from CA3-originating transcallosal fibers to CA1 — also referred to as Schaffer commissurals. In addition, optogenetically induced synaptic input may include the recruitment of other regions through poly-synaptic pathways^10–12,49^, or activation of nearby regions, (e.g., area CA2), due to viral spread to those regions, which themselves may have provided direct or indirect input to CA1. Notably, other studies reporting successful induction of place cells using a similar approach^18^ provided only limited information about success rates (i.e., the distribution of cells non-/responding to the optogenetic stimulation and/or exhibiting a pre-existing/induced/no place field), and therefore, the ratios of induced place cells in the stimulated region vs place cells in other locations or non-place cells is not clear.

In addition, these studies often did not control for the addition of visual landmarks associated with laser light delivery — a phenomenon through which place cells can be induced^29^. By contrast, our experimental approach purposefully comprised cellular recordings also on the days before and after optogenetic manipulations, during which sham light was delivered. In addition to habituating the animals to the laser light, sham light delivery served as an important control for place cell formation evoked by sensory perception. Indeed, these controls allowed us to distinguish place cells induced specifically by optogenetic CA3 stimulation from spontaneously appearing place cells and place cells triggered by perception of an apparent environmental sensory landmark (seeing the laser light), as previously reported^34^. The relatively large fraction of place cells induced by visual perception of the sham light, compared to optogenetically induced place cells, indicates that this is an important confounding factor, which should not be overlooked.

A previous study, using acute silencing of CA3 inputs, already established that CA1 place fields depend on CA3 input and that this manipulation led to a degraded spatial code at the CA1 population level^20^. Our results show that also acute, phasic stimulation of CA3 synaptic input leads to a transient disruption of the CA1 place code. This suggests that any acute disturbance of natural CA3 activity, be it inhibition or stimulation, will lead to a degradation of the spatial code. Despite the ability to acutely control light delivery with high temporal precision, these manipulations do not consider the phase of neuronal oscillations. Thus, activation of CA3 synaptic input may disrupt the spatial code due to a phase mismatch of the optogenetic manipulation with the theta cycle or due to disturbance of theta-gamma coherence^50^. While refined stimulation methods are needed to dissect the role of Schaffer commissures in the regulation of CA1 spatial code with better precision (e.g., by direct, closed-loop stimulation of CA1 axons), our findings provide important insights into of the role of CA3 inputs in the regulation of the dynamics of CA1 spatial representations across days.

Despite changes in the hippocampal code for space, behavioral signatures of navigation, namely running speed and anticipatory licking remained largely unaffected by CA3 optogenetic manipulations. This observation is in line with past reports of place code remapping occurring in parallel of unchanged overt behavior in physiological settings (i.e., without optogenetics)^4−7^. One possible explanation is that unaffected place cells ensure the maintenance of behavior on days where optogenetic manipulations were performed. Alternatively, given that our optogenetic stimulation approach only affected a fraction of all CA3 inputs to CA1, the recurrent connectivity of unaffected CA3^51,52^ and its ability for pattern completion (e.g., in ipsilateral CA3 or more caudal/rostral portions of CA3) could be sufficient to preserve global network activity in a stable, pre-stimulation state.

Taken together, this study underlines the role of CA3 in shaping the dynamics of CA1 spatial coding. Repeated activation of CA3 at a fixed spatial location led to the formation of a small pool of place cells and remapping of the pre-existing spatial code along the entire trajectory of the explored space. More work is necessary to further investigate the mechanisms underlying the formation and (de-)stabilization of spatial representations in individual CA1 cells and how the resulting hippocampal network dynamics contribute to learning and integration of external and internal representations. Future studies could for instance jointly manipulate CA3 activity along with other features involved in spatial coding, including entorhinal cortex, oscillatory activity, neuromodulation and local inhibition to better understand the mechanisms underlying CA1 place cell formation and plasticity.

## METHODS

### Subjects

A total of 4 adult (4 to 9 months of age) C57BL/6J wild-type mice were used in this study. Mice were commercially purchased (*Charles River*, USA) and were housed in pathogen-free conditions at the *University Medical Center Hamburg-Eppendorf* using a light/dark cycle of 12/12 hours respectively. The humidity (∼40%) and temperature (∼20°C) in the room were kept constant. Mice had free access to food, igloos and running wheels in their home cages. All procedures were performed in compliance with German law and according to the guidelines of *Directive 2010/63/EU*. Protocols were approved by the *Behörde für Gesundheit und Verbraucherschutz* of the *City of Hamburg* under the license numbers 32/17 and 33/19.

### Virus injections

The mouse was anesthetized with 2% isoflurane/L of O_2_. After 5 minutes, the mouse was transferred to the surgical area, and anesthesia was maintained with 1.5% isoflurane/L of O_2_. Depth of anesthesia and analgesia was assessed with the paw withdrawal reflex. Buprenorphine (0.1 mg/kg) and carprofen (4 mg/kg) were both injected subcutaneously. The mouse was placed on a heating pad which maintained body temperature at 37°C throughout the surgery. Eye ointment (Vidisic, *Bausch + Lomb*, USA) was applied to prevent eye drying. The fur on the head was carefully trimmed to avoid later contamination of the surgical field. Skin was then disinfected using Betaisodona. The mouse was fixed onto the stereotaxic frame and a 3-4cm midline scalp incision was made close to the injection sites. The skin was pushed to the sides and a bone scarper was used for cleaning (*Fine Science Tools*, Heidelberg, Germany). Two craniotomies were made above the injection sites using a dental drill (*Foredom*, USA). First, 300nL of an AAV2/9-CaMKII-ChrimsonR-mScarlet-KV2.1 (1.10^13^ vg/mL, 124651, *AddGene*, USA) viral suspension was injected in left CA3 (−2.0mm AP, −2.3mm ML, −2.5mm DV relative to bregma) using a custom-made air-pressure driven injection system. Then, 500nL of AAV2/9-Syn-jGCaMP8m (4.10^12^ vg/mL, 162375, *AddGene*, USA) were injected in right CA1 (−2.0mm AP, +1.5mm ML, −1.3mm DV relative to bregma). The scalp was then sutured. Mice were removed from anesthesia and let to recover in a dedicated cage half placed on a heating blanket. For three days after surgery, mice were provided with Meloxicam mixed with soft food.

### Hippocampal window surgery

After a minimum of two weeks post virus injections, a second surgery for hippocampal window implantation was performed following the same steps described above. A first craniotomy was drilled above left CA3 (−2.0mm AP, −3.5mm ML, 35° angle towards the left) and an optic fiber (1.25mm ferrule diameter, 0.22 NA, 1.3 ± 0.1mm length, *Doric Lenses*, Canada) was inserted and glued to the skull. A circular 3mm diameter bone piece, centered around right CA1 injection site, was carefully removed using a trephine (ISO 020, *MW Dental*, USA). The dura and somatosensory cortex above the hippocampus were carefully aspirated until the white matter tracts of the corpus callosum became visible. A drop of transparent Kwik-Sil was applied in order to reduce motion-related artefacts in the recordings of behaving mice^53^. Sterile PBS was used to wash the craniotomy all along, and a custom-made hippocampal imaging window was inserted. The cranial window consisted of a hollow glass cylinder glued to a No. 1 coverslip using UV-curable glass glue (Norland NOA61, *Cranbury*, USA). The imaging window and a head plate (*Luigs & Neumann*, Germany) were attached to the skull with cyanoacrylate gel (SuperGel, *UHU*, Germany). Dental cement (Super Bond C&B, *Sun Medical*, USA) was then applied to seal the implant. Animals were provided with post-operative care and let to recover for a minimum of two weeks before behavioral training was initiated.

### Two-photon imaging

After the mouse was head-fixed on a linear treadmill (*Luigs and Neumann*, Germany), the hippocampal cranial window was centered under the two-photon microscope using epifluorescence, and for each mouse, a field of view was chosen on the first recording session and returned to at each new session. Single planes (512 × 512 pixels) were acquired using a Ti:Sa laser (930nm, 30-100mW, Chameleon Vision-S, *Coherent*, USA) and a 16× water immersion objective (CFI 16×, 0.80NA, 3mm WD, *Nikon*, The Netherlands) at a 30Hz acquisition rate using ScanImage (v2017b; *MBF Bioscience*, USA).

### Optogenetic stimulation

The optogenetic stimulation laser (LuxX 633-100, LightHUB*, Omicron*, Germany) was connected to the fiber optic implant using a custom-made ceramic sleeve allowing intra- or extracranial light delivery (for sham stimulation). Optogenetic stimulation consisted of 1s continuous illumination of 633nm light at a power of 10mW at the brain end. Both sham and optogenetic light delivery was triggered in laps #21 to #30 at a fixed, predefined location on the treadmill belt. Optogenetic stimulation was delivered at the same treadmill location for all mice. Sham stimulations occurred at one of 4 equally spaced, predefined locations which was randomly allocated at each session for each mouse.

### Position monitoring

Mice ran on a 2-meter-long treadmill belt composed of 5 identical length sections with distinct tactile cues. A white band placed under the belt signaled lap reset by reflecting an infrared light beam. Mouse position was monitored using a rotary encoder (1024 positions, E6B2-CWZ6GH, *Yumo*, China) attached to a 3D-printed custom-engineered wheel under the belt, and connected to a rotary encoder module (1034, *Sanworks*, USA) sending angular position through an analog channel to the master NI board co-registering behavioral events with 2p frame acquisition (at a sampling rate of 1.5kHz) and optogenetic stimulation light triggering.

### Lick detection

A high-speed camera (600fps, DMK 33UX252, *The Imaging Source*, Germany) streamed a lateral view of the infrared-illuminated mouse. The camera stream was processed online using a custom-made *Bonsai* workflow^54^. In short, a region of interest (ROI) was manually drawn around the tip of the lick tube and licks were detected whenever the sum of ROI pixel values crossed a manually predefined threshold.

### Task logic

Task events were monitored and triggered using the *Bpod* finite state machine (1035, *Sanworks*, USA). *Bonsai* communicated lick occurrence to *Bpod* (through a local TCP protocol), which triggered a digital signal to the master NI board (*National Instruments*, USA). Whenever a lick occurred in the reward zone, *Bpod* also triggered water reward delivery (4μL) by opening a solenoid valve (LHDA1231115H, *Lee Company*, USA) connected to a port interface board (1004, *Sanworks*, USA).

### Shaping protocol

Mice were gradually habituated to the experimenter, the experimental setup and to head fixation, in that order. The weight of each mouse was monitored daily and maintained at ≥ 85% of their initial weight. On the first day following water restriction, mice learned to collect water reward drops from the lick tube: first, water reward drops were delivered by the experimenter; next, mice received water whenever they licked the reward spout. In the second phase of training, typically starting on the second day and lasting for 2-3 days, mice had to run on the treadmill to obtain an automatically provided (no lick necessary) water reward drop at the reward zone. On the last phase of training, mice had to lick in the reward zone to obtain the water reward drop. Task expertise was identified by uninterrupted running for a minimum of 50 laps and selective licking in the reward zone. Throughout training, sham light was delivered (see above) for habituation purposes and to prevent behavioral alterations by optogenetic stimulation in experimental sessions. In total, shaping lasted for 5 to 10 days.

### Histology

Mice were injected with a lethal dose of ketamine and xylazine before being transcardially perfused with 1x PBS followed by 4% paraformaldehyde (PFA). Brains were removed, stored in 4% PFA and sliced using a vibratome (60μm thickness, VT1000S, *Leica*). Brain slices were mounted on microscope slides using an aqueous mounting medium (FluoroMount, *ThermoFisher*, USA) and coverslips (1871, *Carl Roth*, Germany). Full overview images were acquired with an epifluorescent microscope (AxioObserver, *Zeiss*, Germany) equipped with a 10x objective (Plan-Apochromat, *Zeiss*, Germany). Detailed views of region of interests were acquired with a confocal microscope (LSM 900, *Zeiss*, Germany) equipped with a 20x objective (Plan-Apochromat, *Zeiss*, Germany).

### Preprocessing of behavioral signals

Digital signals conveying licks, reward zone, and optogenetic triggers were binarized and down-sampled to the sampling rate of the 2p-microscope (30Hz). Position (0 to 2m) was estimated by quantizing (5cm-long bins, total of 40 bins) the corresponding analog signal (−4.5 to 4.5V) and down-sampling it to the sampling rate of the 2p-microscope. Speed was then taken as the first-order derivative of animal position.

### Preprocessing of calcium activity

For each mouse, all recordings were provided to *Suite2p* (version 0.13.0) to correct for motion-related plane distortion and to co-register frames across sessions. Individual cells were identified and segmented based on morphology only using *CellPose*^55^. Neurons were manually sorted and labeled when not automatically identified. Time points with speed lower than 10cm⋅s^−1^ were excluded from further analysis. To correct for bleaching of the calcium indicator, time-resolved baseline correction was applied separately for each neuron: first, surrounding neuropil fluorescence (*F*_neuropil_) was removed from the cell fluorescence (*F*_cell_) as follows: *F* = *F*_cell_ − 0.7 × *F*_neuropil_ (following past work^46−48^); second, Δ*F*/*F*_0_ = (*F − F*_0_) / *F*_0_ was computed, where *F*_0_ corresponds to the 30th-percentile on a 3s moving window. To reduce noise, the Δ*F*/*F*_0_ trace was further smoothed using a 166ms (5 bins) moving boxcar.

### Identification of responding neurons

For each neuron, the baseline corrected (baseline = 2s before stimulation onset) trial average (10 trials) calcium response locked to optogenetic stimulation onset was computed. A neuron was classified as responding if more than 50% of the time points belonging to the optogenetic stimulation window (0-1s) were greater than 97.5% of the time shuffled distribution of Δ*F*/*F*_0_ values (10,000 shuffles, *P* < 0.05, two-sided).

### Motion correction

Before identifying place cells, we aimed to further correct for out-of-plane motion displacement. We based our approach on the previously proposed idea that z-motion would produce an equal number of positive- and negative-going (false) calcium transients^59^. In our analysis, positive- and negative-going calcium transients corresponded to time-points with a Δ*F*/*F*_0_ respectively above/below *k* × MAD ± median(Δ*F*/*F*_0_) thresholds. The hyperparameter *k* was defined independently for each cell such that the lower error rate was smaller or equal to 5% (i.e., *N*(negative transients) / *N*(positive transients) ≤ 0.05). Whenever Δ*F*/*F*_0_ was below *k* × MAD + median(Δ*F*/*F*_0_), Δ*F*/*F*_0_ was then set to zero for further analyses.

### Identification of place cells

For each cell, a tuning curve was computed by averaging Δ*F*/*F*_0_ separately for each quantized position along the treadmill. A cell was qualified as a ‘place cell’ if the following criteria were met: (i) significant points (see below) on the tuning curve spanned a window of 10 to 75cm (2 to 15 bins), and (ii) tuning curves fulfilling criterion #1 were found in at least 30% of the following laps. A significant point on the tuning curve is a position bin in which the average Δ*F*/*F*_0_ was greater than 97.5% of the null distribution of Δ*F*/*F*_0_ obtained after shuffling animal position in time (offset of at least 5s, 10,000 permutations).

### Maps of place tuning

To avoid exaggerating the diagonal pattern in place cell × position activity maps, the ranks of place cells were cross validated. To this end, the sub-rank of each place cell was obtained by finding the peak of the tuning curve estimated separately on odd vs. even laps. Place cells were then ranked according to the average of odd- and even-based sub-ranks. The tuning curves estimating separately on odd vs. even laps were then averaged to yield a cross-validated rank and tuning curve for each cell. For comparison of place cell tuning across days, the cross validated cell rank obtained on a given day was applied to the other days.

### First lap response

The lap at which induced place cells show the first sign of place tuning was determined. This lap corresponded to the lap in which the average Δ*F*/*F*_0_ of the induced place field (determined on the average across laps) was greater than 97.5% of the null distribution of Δ*F*/*F*_0_ obtained after shuffling animal position in time.

### Bayesian decoder of position

The position of the animal was decoded at each time-point using the activity of all place cells, as follows^6,33,60–62^:

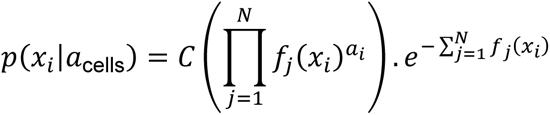

where *f*_*j*_(*x*_*i*_) is the template activity of place cell *j* at position *x*_*i*_, *a*_*i*_ is the recorded activity in the current time-point, and *C* is a normalization constant (ensuring the distribution integrates to 1) chosen to implement a uniform prior over positions. The decoded position was taken as the maximum a posteriori and the decoding error was measured as the absolute difference between the decoded and the true position. Depending on the question of interest (see main text), the decoder was trained (i.e., ***f***(***x***) estimated) and tested on the same or different sets of laps. To provide a fair comparison when the decoder was trained and tested on the same set of laps, decoder training and testing were systematically 2-fold cross validated by analyzing odd and even laps separately: (i) training on odd laps and testing on even laps, (ii) training on even laps and testing on odd laps, and (iii) averaging decoding error in each of the 2 folds. For comparison between pre and post laps, the number of laps was kept equal:(20 laps) by discarding late post laps.

### Place field shift

To quantify the shift in place field between pre and post laps, we computed two place turning curves for each place cell exhibiting a place preference in pre laps. Place field shift was then measured as the absolute distance between the position (in cm) of the peak of each of the two place tuning curves.

### Place field precision

To detect a change in the precision (i.e., inverse variance) of place fields, the place tuning curve was (i) reshaped and expressed as a function of the absolute distance to its peak and (ii) normalized by the peak fluorescence. The difference between the peak-aligned place tuning curves in post vs. pre laps gives a measure of the amount of change in place field precision.

### Statistics

Statistical analyses were performed using custom-made Python and R scripts. Whenever specified, permutation tests were performed to generate a null distribution. To do so, 10,000 random data shuffling were performed. When the number of all possible permutations was smaller than 10,000, all possible permutations were listed and used. Significance thresholds correspond to 2.5% and 97.5% percentiles of the resulting null distribution (*P* < 0.05, two-sided). The *P*-value was obtained by computing the proportion of random samples with values at least larger or smaller than the observed test statistic (e.g., a difference between conditions).

## ACKNOWLEDGEMENTS

We thank Michael D. Adoff and Daniel A. Dombeck for advice on cranial window surgery as well as Stefan Schillemeit and Kathrin Sauter for administrative and technical support throughout the project. MM is supported by the *Alexander von Humboldt Foundation* and by the *Fondation Bettencourt-Schueller*. This work was funded by the Deutsche Forschungsgemeinschaft (DFG, German Research Foundation) SFB 936 - 178316478 − B8 and FOR2419 − 278170285 − P6.

## DATA AND CODE AVAILABILITY

The code and datasets are available upon request.

## AUTHOR CONTRIBUTIONS

CR, MM and JSW designed the experiments. CR performed the experiments. CR and MM analyzed the data and made the figures. CR and MM wrote the manuscript, together with JSW. JSW and MM secured funding.

## DECLARATION OF INTERESTS

The authors declare no competing interests.

CR is currently a Poitras Center Fellow at the *McGovern Institute*, *MIT*, Cambridge, MA, USA.

MM is currently a Marie Skłodowska-Curie Postdoctoral Fellow at the *Department of Psychological and Brain Sciences*, *Boston University*, Boston, MA, USA.

## SUPPLEMENTARY FIGURES

**Supplementary Figure 1.**
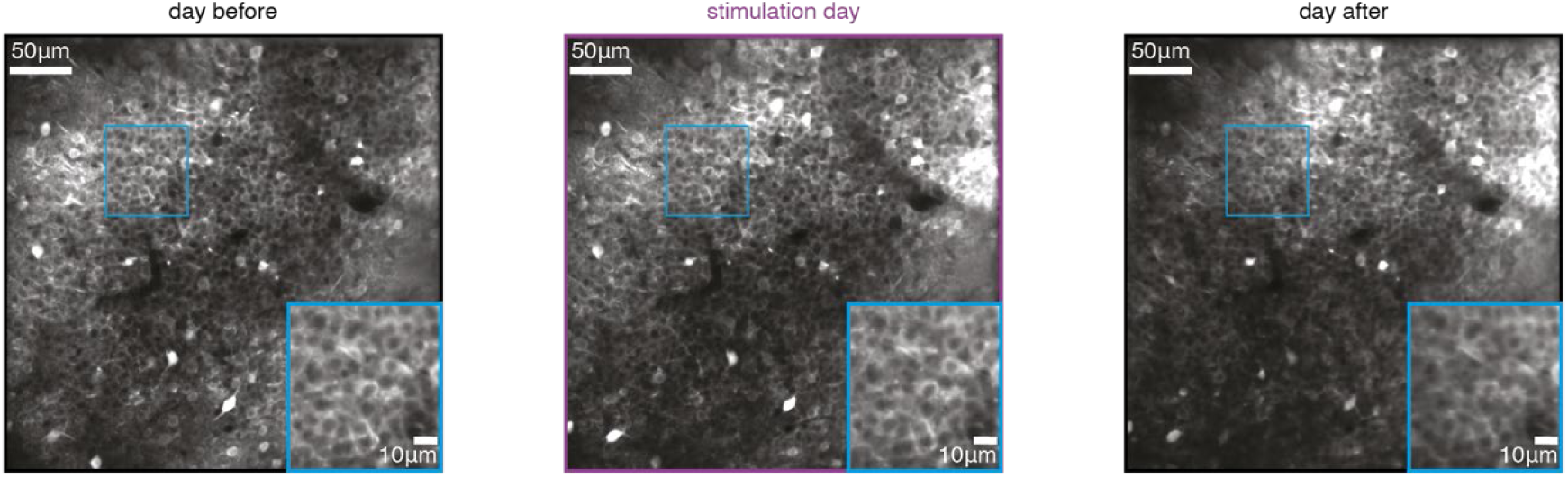
Example field of view across days. Field of view of an example mouse on day before (left), stim day (middle) and day after (right). Insets represent a magnified view of the region indicated by the blue square across days.

**Supplementary Figure 2.**
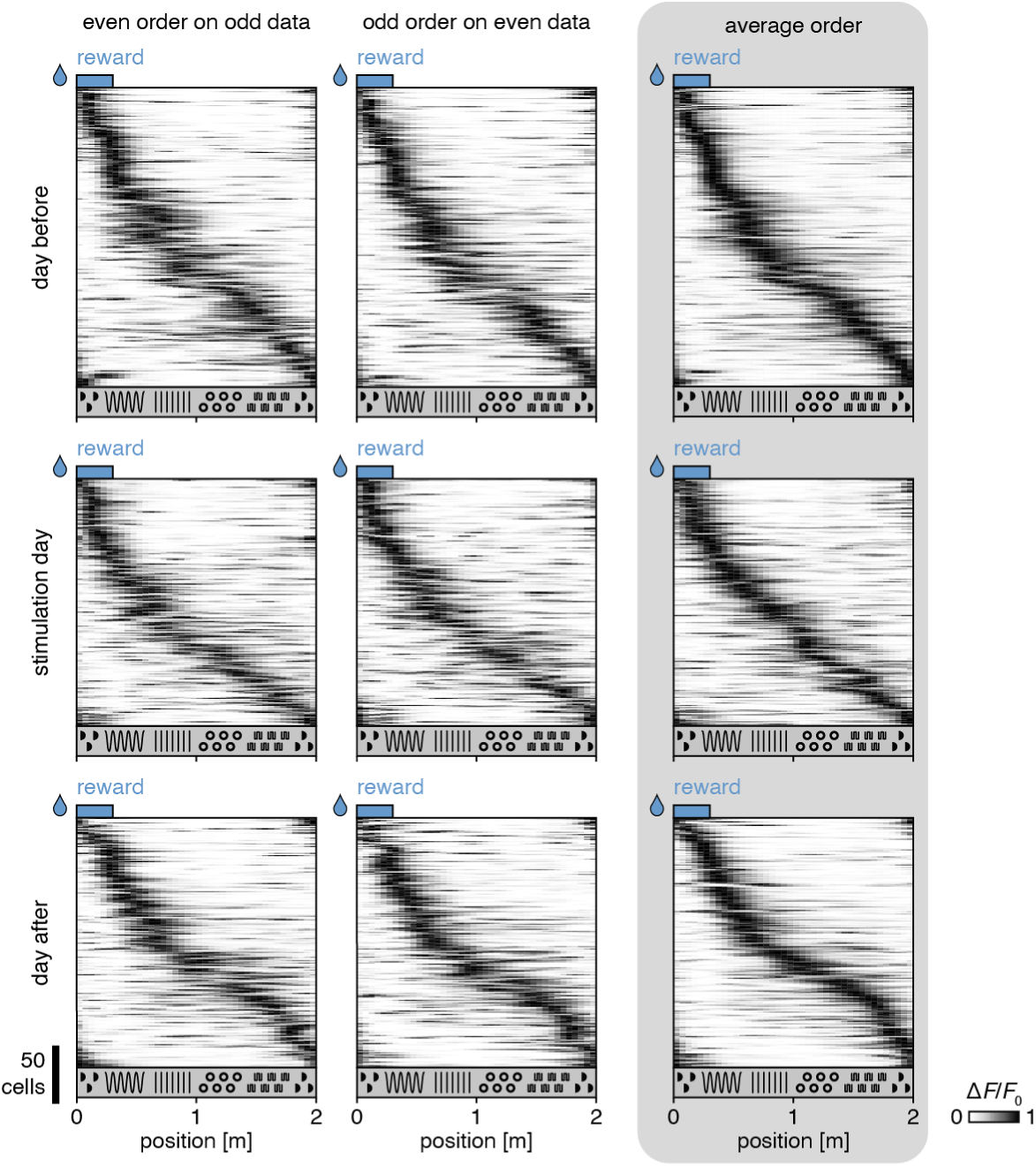
Cross-validation of place cell maps. Place cells representing the day before (top row), stimulation day (middle row) and day after (bottom row) the optogenetic stimulation day are ranked based on the rank obtained by averaging activity on even laps and odd laps separately (right column). The ranks of average activity in even laps are applied to activity in odd laps (left column) and vice versa (middle column).

**Supplementary Figure 3.**
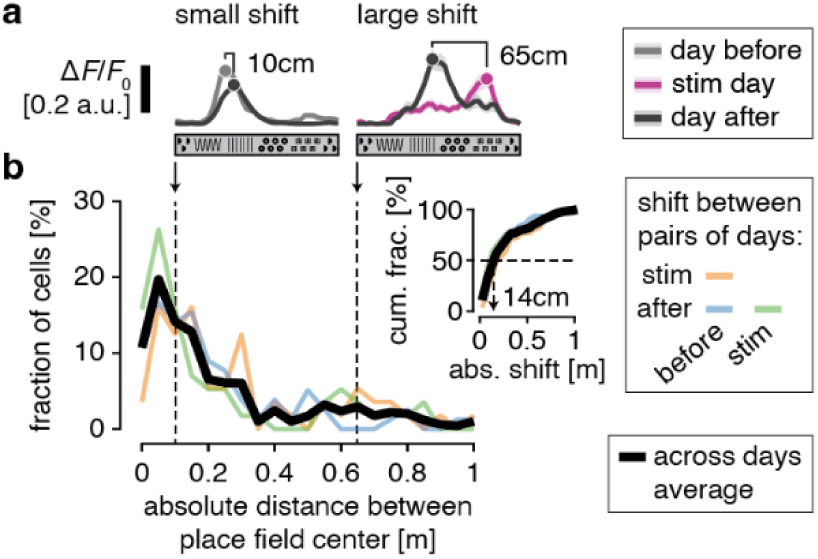
Persistent place cells encode close-by locations across days. **a**, Two example persistent place cells with a small (10cm) and large (65cm) shift in place field center on two different days. **b**, Fraction of place cells as a function of their shift of the place field center from one day to another (before vs. stimulation day, before vs. after stimulation day, stimulation day vs. after), and after averaging across all 3 combinations of days (black). Inset shows the corresponding cumulative distribution. Half of persistent place cells have a shift smaller than or equal to 14cm.

